# Beneficial effects of novel strains of Aureobasidium pullulans produced 1,3-1,6 β-glucans on non-esterified fatty acid levels in diabetic KKAy mice

**DOI:** 10.1101/2021.07.22.453362

**Authors:** Nobunao Ikewaki, Takashi Onaka, Yasunori Ikeue, Mitsuru Nagataki, Gene Kurosawa, Vidyasagar Devaprasad Dedeepiya, Mathaiyan Rajmohan, Suryaprakash Vaddi, Rajappa Senthilkumar, Senthilkumar Preethy, Samuel JK Abraham

## Abstract

Obesity, metabolic syndrome, associated lipotoxicity and its cascade of events contribute to the majority of the burden related to non-communicable diseases globally. Preventive lifestyle changes aside, several beneficial effects have been reported in type II diabetes mellitus and dyslipidaemia patients with biological response modifier glucans (BRMG) produced as an exopolysaccharide by *Aureobasidium pullulans*. In this study, we compared two strains (AFO-202 and N-163) that produce beta glucans in alleviating lipotoxicity. This study was performed in obese diabetic mice model of KK-Ay mice, in four groups with six subjects in each group - Group 1: sacrificed on Day 0 for baseline values; Group 2: control (drinking water); Group 3: AFO-202 beta glucan—200 mg/kg/day; Group 4: N-163 beta glucan—300 mg/kg/day. The animals in groups 2–4 had the test solutions administered by gavage once daily for 28 consecutive days. Biochemical analyses were conducted of blood glucose, triglycerides, total cholesterol, LDL cholesterol, HDL cholesterol and non-esterified fatty acids (NEFA). Group 4 (N-163) had the lowest NEFA levels, as compared to the other groups, and marginally decreased triglyceride levels. The groups had no significant differences in blood glucose, HbA1c, triglycerides, or LDL and HDL cholesterol. N-163 produced by *A. pullulans* decreased NEFA in a diabetic mice model in 28 days. These results, although modest, warrant further in-depth research into lipotoxicity and associated inflammatory cascades in both healthy and disease affected subjects to develop novel strategies for prevention and management.

## Introduction

Fatty acids are a major component of lipids and are an important source of fat fuel for the body. The non-esterified fatty acids (NEFA) of free fatty acids (FFA) are the circulating form of fatty acids in the plasma [1]. FFA are an important link between obesity, insulin resistance and inflammation and the development of diabetes, hypertension, dyslipidemia, coagulative disorders and cardiac diseases [2]. Lipotoxicity due to the destructive effects of excess fat accumulation impairs the function of several metabolic pathways [3]. In particular, chronic excessive levels of plasma FFA, leading to insulin resistance, influences the synthesis of hepatic triglycerides. The hepatic fatty acid metabolism, which is closely linked to inflammation, leads to hepatic steatosis (non-alcoholic steatohepatitis [NASH]), progressing to fatty liver disease, cirrhosis and cancer [3]. Therefore, modulating and normalizing NEFA can considered as a key target to alleviate the effects of the entire cascade of lipotoxicity [4] and the dysregulation of lipid metabolism described above. Both statin and non-statin pharmacological agents have been studied for their effects on NEFA, with modest outcomes [5].

Beta glucans are polysaccharides with many beneficial effects to ameliorate glucose metabolic disorders such as diabetes and dyslipidaemia [6], in addition to enhancing immunities for fighting viral infections and cancer [7]. The effects have been reported of 1,3-1,6 beta glucan derived from the AFO-202 strain of the black yeast *Aureobasidium pullulans* (Figure 1A) in individuals with Type II diabetes [8] and dyslipidaemia [9]. N-163 is another strain of *A. pullulans* which produces a novel variant of 1,3-1,6 beta glucan (Figure 1B). In vitro studies of N-163 beta glucan have shown its positive influence on decreasing inflammatory cytokines (data unpublished). For this study, we have evaluated the effects of beta glucans derived from AFO-202 and N-163 *A. pullulans* in KK-Ay mice, as obese diabetic mice models. KK-Ay mice are applied as animal models for research on obesity and metabolic disorders, and they were originally developed by crossing KK mice with yellow obese mice (Ay mice) [10].

**Figure 1:**
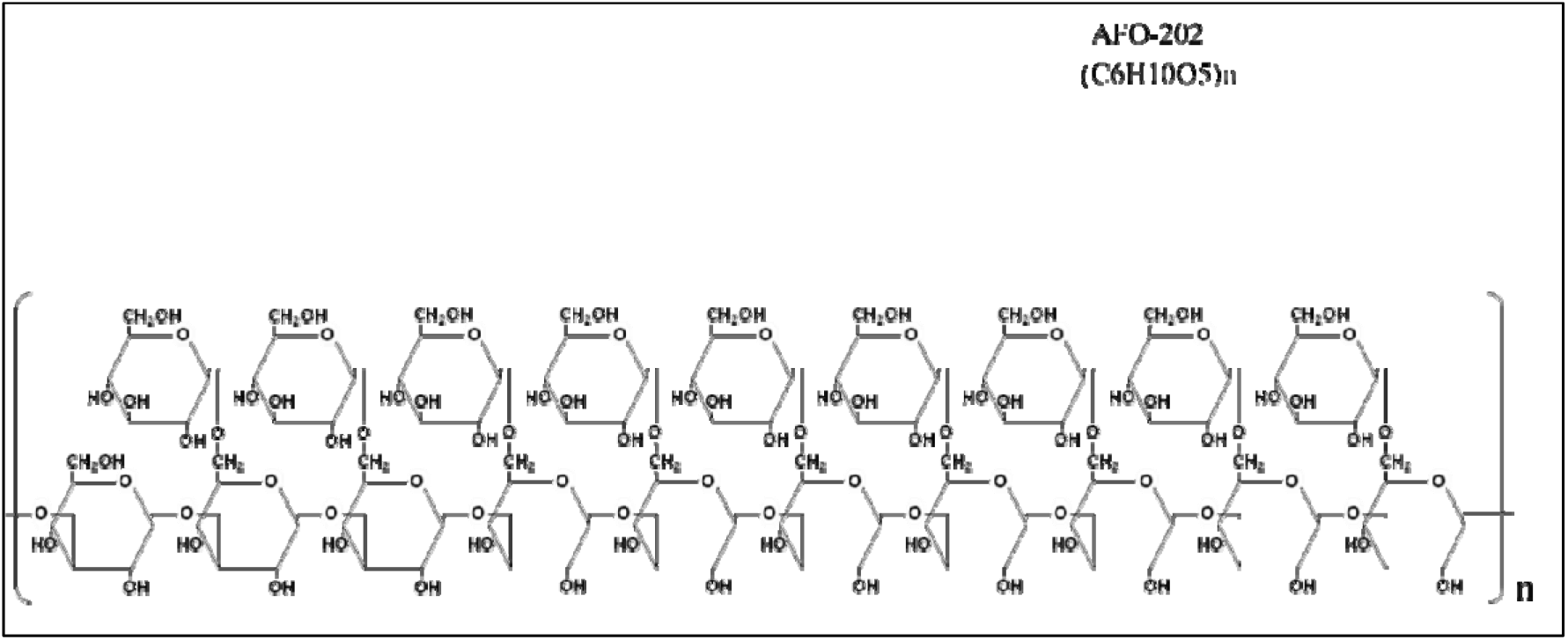
Structure of beta glucan derived from the AFO-202 strain with the chemical formula (C_6_H_10_O_5_)n

## Methods

### AFO-202 and N-163 beta glucans

*A. Pullulans* is a harmless, naturally occurring black yeast that originally was isolated from soil. Common cultures base on potato dextrose agar (PDA) and potato dextrose broth (PDB) were used for the initial lab-level cultures and later, when scaled up for large cultures using each specific medium so that the AFO-202 and N-163 strains would produce β-glucan as an exo-polysaccharide with a chemical structure of (C_6_H_10_O_5_)n. For the AFO-202 culture, rice bran was used as a nitrogen source along with a natural medium consisting mainly of rice bran, ascorbic acid and glucose. The N-163 strain was cultured with nitrates instead of rice bran and a synthetic medium consisting of thiamine nitrate, yeast extract and glucose. The storage and incubation temperatures for both strains were around 25 °C. The incubation period was five days for the N-163 strain and six days for the AFO-202 strain. The produce, in gel form, then was heat-sterilised to yield a consumable product. The pH of both products was 5.0 ± 1.0. The produce of the N-163 strain was more viscous and formed more threads, as compared to the produce from the AFO-202 strain. This difference may be due to the difference in the structural formula of β-glucans (Figure 1, 2).

**Figure 2:**
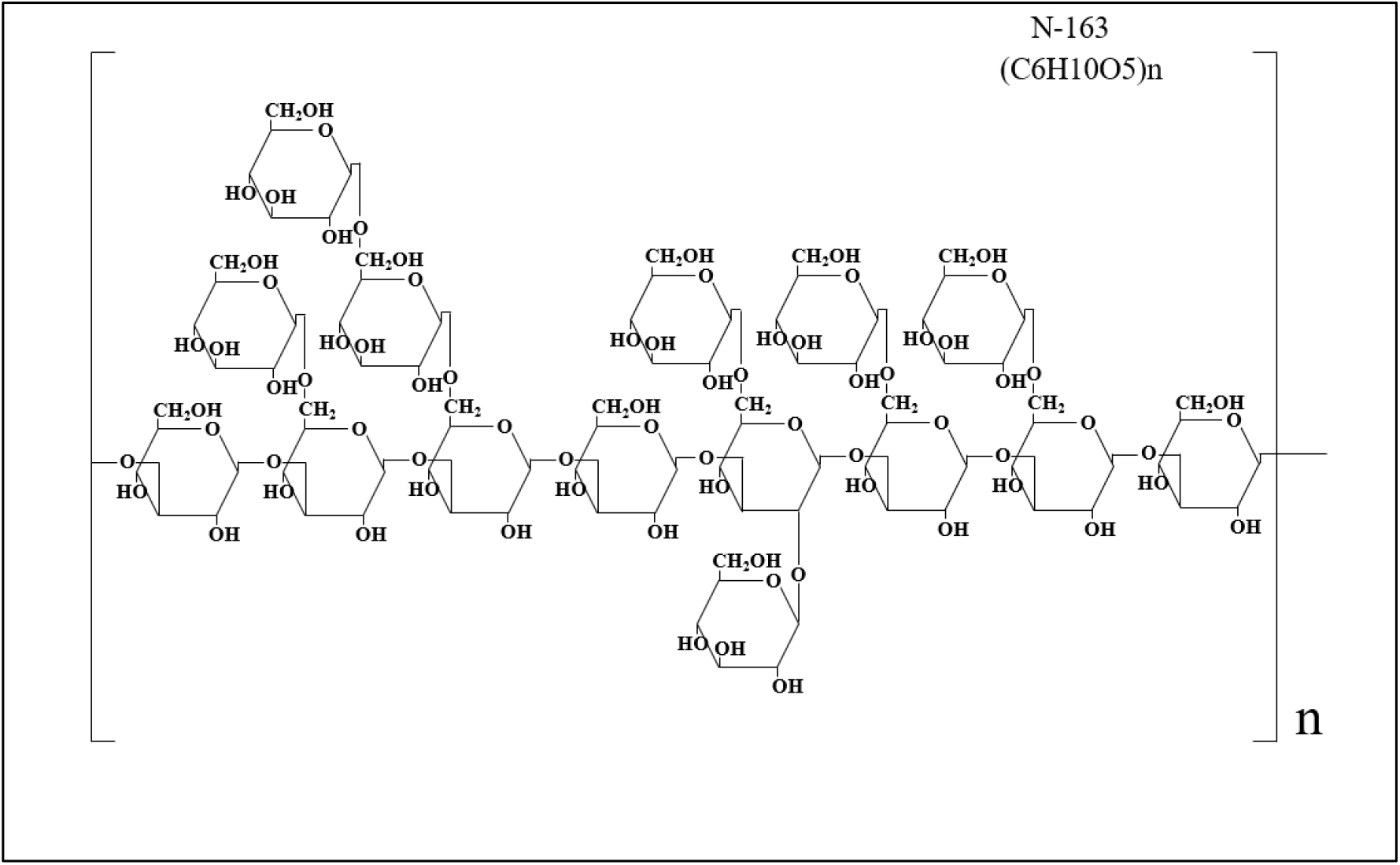
Structure of strain beta glucan derived from N-163 with the chemical formula C_6_H_10_O_5_)n

### Animal study

Protocol approval was obtained from the Ethics Committee of Toya Laboratory, HOKUDO Co. (ref. no. HKD47047). The study was conducted in accordance with the HOKUDO Animal Experiment Regulations following the Act on Welfare and Management of Animals (Ministry of the Environment, Japan, Act No. 105 of October 1, 1973), standards relating to the care and management of laboratory animals and relief of pain (Notice No. 88 of the Ministry of the Environment, Japan, April 28, 2006) and the guidelines for proper conduct of animal experiments (Science Council of Japan, June 1, 2006).

Healthy 6-week -old male KK-Ay/TaJcL mice (Nippon Clare Co., Japan) were purchased for the study. The animals were acclimatized for 3 weeks from their date of arrival. Healthy animals with no abnormalities during the acclimatization period were divided into four groups (six males per group) using a weight-stratified randomization method so that the average weights of each group would be as uniform as possible. The animals were kept in a rearing environment with room temperature of 23 ± 2 °C (acceptable limit range: 20–26 °C), a relative humidity of 55 ± 10% (acceptable limit range: 30–70%) and 12 hours each of light and dark (light hours: 7:00 a.m. to 7:00 p.m.). The mice were housed in microbarrier cages made of polysulfone (external dimensions: W196 mm × D306 mm × H166 mm) with bedding chips (Dohoh Rika Sangyo Co., Ltd.). One mouse was kept in each cage. The cage and bedding were changed at least once a week. The animals were fed a solid feed (CE-2, Feed One Co., Ltd.), which was manufactured within the past year. The solution used for all the groups was groundwater that had been sterilized by adding sodium hypochlorite to achieve a residual chlorine level of 0.3-–.4 mg/L using a facility water sterilizer (MJ25SR, Kawamoto Manufacturing Co., Ltd.). The water bottles were changed at least twice a week.

The groups and doses of the test substances were as follows.

Group 1 (n = 6): Sacrificed on Day 0 for baseline values

Group 2 (n = 6): Control (solvent—drinking water)

Group 3 (n = 6): AFO-202 beta glucan—200 mg/kg/day; 20 mg/ml concentration in solution

Group 4 (n = 6): N-163 beta glucan—300 mg/kg/day; 30 mg/ml concentration in solution

The dose of each test substance was determined to be the same as the expected daily intake dose for humans. In other words, the estimated daily human intake of each test substance is 10 g in gel form of AFO-202 beta-glucan (5 mg of active ingredient of beta-glucan per gm) and 15 g in gel form of N-163 beta-glucan (6 mg of active ingredient of beta-glucan per gm).

Because the test substance is a food material, oral administration via gavage was selected, which is commonly used for oral administration in rodents. The administration period was 28 days. The animals were forcibly administered orally into the stomach using a gastric tube (KN-348, oral administration needle, Natsume Corporation) and a disposable syringe (Terumo Corporation) once daily for 28 consecutive days (between 08:00 and 15:00).

The animals in Group 1 were weighed using an electronic balance (FX-1200I, A&D Co., Ltd.) on the day before the start of treatment (Day 0). Afterward, all of the animals underwent laparotomy under isoflurane anaesthesia (isoflurane, Fujifilm Wako Pure Chemical Co., Ltd.), and blood was collected from the aorta. The blood was divided into two portions, one of which was heparinized, and the whole blood was frozen for HbA1c measurement. The other portion was centrifuged to obtain serum, which then was frozen and stored to measure blood glucose, triglyceride, total cholesterol, LDL cholesterol, HDL cholesterol and free fatty acid measurements. The centrifugation conditions were 3500 rpm at 4 °C for 15 minutes.

The general condition of all subjects in Groups 2 and 3 was observed at least once a day from the day of administration (Day 1) to the day of autopsy. Body weight was measured before dosing (between 08:00 and 13:00) on days 1, 7, 14, 21 and 28 (last dosing day) for all subjects in Groups 2, 3 and 4. On the day of necropsy, the body weight of each mouse was measured before fasting. Body weight was measured using an electronic balance (FX-1200I, A&D Co., Ltd.).

The average food intake per day was calculated from the food intake over 3 days. To measure the acclimation period, the amount of food fed on Day 18 and the amount of food remaining on Day 21 were measured for all cases. For the acclimation period, the feed intake was measured on Days 3, 10, 17 and 24, and the residual feed intake was measured on Days 6, 13, 20 and 27. From these data, the mean daily food intake was calculated at doses 3–6, 10–13, 17–20 and 24–27.

Blood glucose levels were measured by a blood glucose monitoring device (Freestyle Freedom Lite, Nipro Corporation) using whole blood obtained from the tail vein on the day before the start of dosing (Dosing Day 0). The blood glucose level was also measured using whole blood obtained from the tail vein in the same way before the afternoon (13:00–15:00) after 14 days of administration.

The blood glucose, triglycerides, total cholesterol, LDL cholesterol, HDL cholesterol and NEFA tests were conducted using the refrigerated blood and frozen serum sampled before the start of dosing (Day 0) and the frozen blood and frozen serum sampled on the day following the last day of dosing (Day 29). The tests were outsourced to Nagahama Life Science Laboratory, Japan.

We calculated the mean body weight and standard deviation for each group days 1, 7, 14, 21 and 28; average daily food intake during the acclimation period; average daily food intake on days 3–6, 10–13, 17–20 and 24–27; blood glucose levels over time; HbA1c on the day after the last dose; blood glucose in serum; and triglycerides. Total means and standard deviations were calculated for each group for total cholesterol, LDL cholesterol, HDL cholesterol and free fatty acids. Furthermore, the Bartlett test was used to test for equality of variance using Excel statistics (Social Information Service Co., Ltd.). Equal variances were analysed using one-way ANOVAs, and unequal variances were analysed using Kruskal–Wallis tests. When a significant difference was found in a one-way ANOVA, Dunnett’s multiple comparison test was used to compare the means with the control group. When significant differences were found using a Kruskal–Wallis test, the means were compared with the control group using Dunnett’s nonparametric multiple comparison test. The significance level was set at less than 0.05.

## Results

The mean weight of the animals in Group 1 was 41.47 g. Their average daily food intake was 6.71 g/day during the acclimation period of 18 to 21 days. Their blood glucose was 730 mg/dL, HbA1c was 6.89%, triglycerides were 663 mg/dL, total cholesterol was 164 mg/dL, LDL cholesterol was 7 mg/dL, HDL cholesterol was 96 mg/dL and free fatty acids were 1113 μEq/L (Table 1).

**Table 1:**
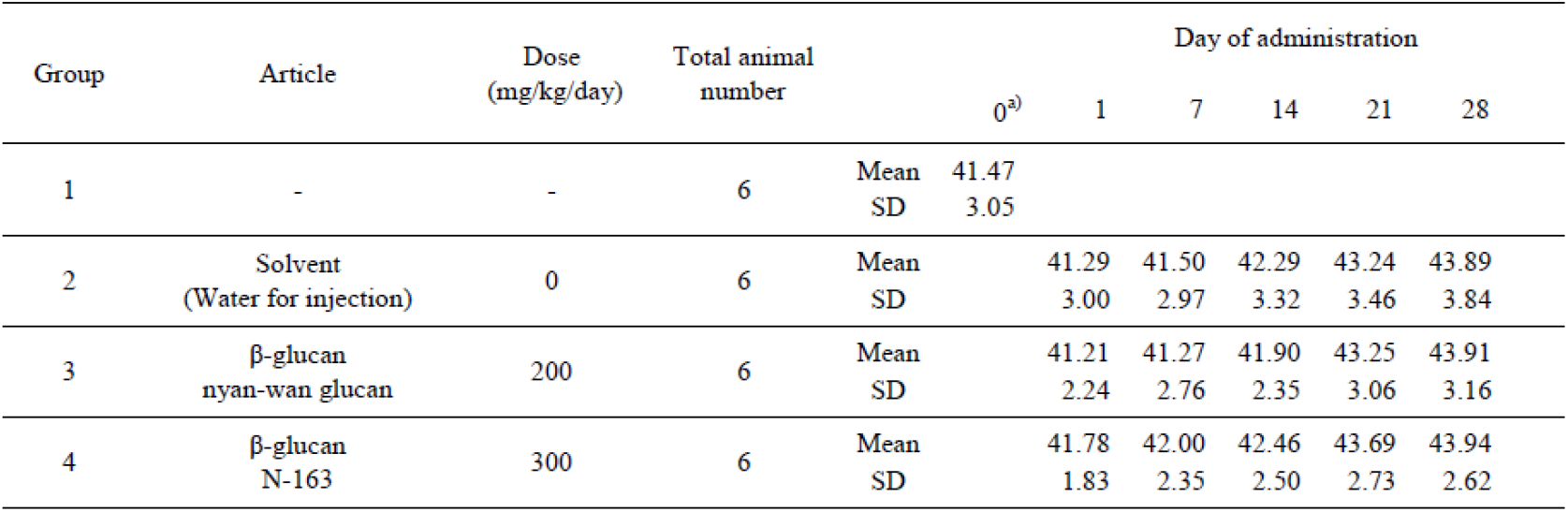
Body weight changes in male KK-Ay mice administered test solutions orally for 28 days

There were no significant differences in body weight or food intake among Groups 2, 3 and 4. None of the animals showed any dose-related abnormalities.

After 28 days, the average blood glucose level was 592 mg/dL in Group 2, 615 mg/dL in Group 3 and 599 mg/dL in Group 4. The differences were not significant. At the end of the treatment period, the HbA1c of each test substance and the control ranged between 7.85% and 8.35%. There was no significant difference in blood glucose, HbA1C, total, or LDL or HDL cholesterol between the groups (Table 2) at Days 14 and 28 of treatment.

**Table 2:** Blood chemical findings in male KK-Ay mice administered test solutions orally for 28 days

| Group | Article | Dose<br>(mg/kg/day) | Total<br>animal<br>number | | Glucose<br>(mg/dL) | HbA1c<br>(%) | Triglyceride<br>(mg/dL) | Total-<br>cholesterol<br>(mg/dL) | LDL-Cholesterol<br>(mg/dL) | HDL-Cholesterol<br>(mg/dL) | NEFA<br>(μEq/L) |
| --- | --- | --- | --- | --- | --- | --- | --- | --- | --- | --- | --- |
| 1<sup>a)</sup> | - | - | 6 | Mean<br>SD | 730<br>63 | 6.89<br>0.37 | 663<br>121 | 164<br>19 | 7<br>1 | 96<br>9 | 1,113<br>103 |
| 2 | Solvent<br>(Water for<br>injection) | 0 | 6 | Mean<br>SD | 592<br>86 | 8.21<br>0.67 | 521<br>183 | 179<br>26 | 9<br>3 | 111<br>11 | 1,888<br>830 |
| 3 | β-glucan<br>nyan-wan<br>glucan | 200 | 6 | Mean<br>SD | 615<br>95 | 8.17<br>0.70 | 499<br>142 | 181<br>24 | 10<br>2 | 114<br>13 | 1,886<br>456 |
| 4 | β-glucan<br>N-163 | 300 | 6 | Mean<br>SD | 599<br>96 | 8.26<br>0.41 | 491<br>114 | 179<br>17 | 10<br>2 | 110<br>11 | 1,691<br>536 |

NEFA levels were lowest in Group 4 (N-163) (mean ± SD 1691 ± 536 mg/dL), as compared to Groups 3 (AFO-202) (1886 ± 456 mg/dL) and 2 (control) (1888 ± 830 mg/dL) (Figure 3). Similarly, triglyceride levels were the lowest in Group 4 (N-163) (491± 114 mg/dL), as compared to 499 ± 142 mg/dL in Group 3 (AFO-202) and 521 ± 183 mg/dL in Group 2 (control) (Figure 4).

**Figure 3:**
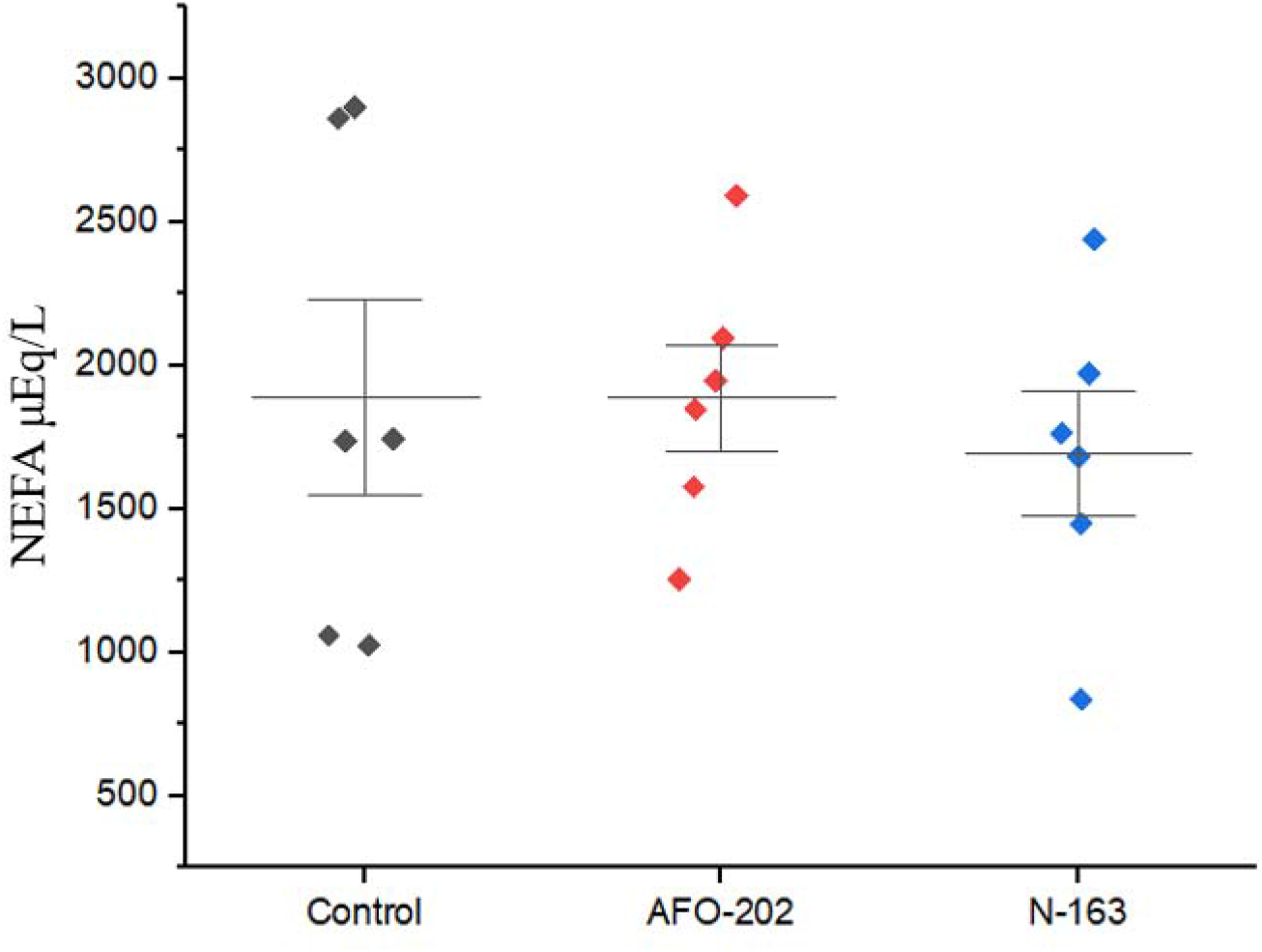
NEFA levels in male KK-Ay mice administered solution orally for 28 days, showing better reduction in Group 3 (N-163 beta glucan) compared to Groups 2 (AFO-202 beta glucan) and 1 (control)

**Figure 4:**
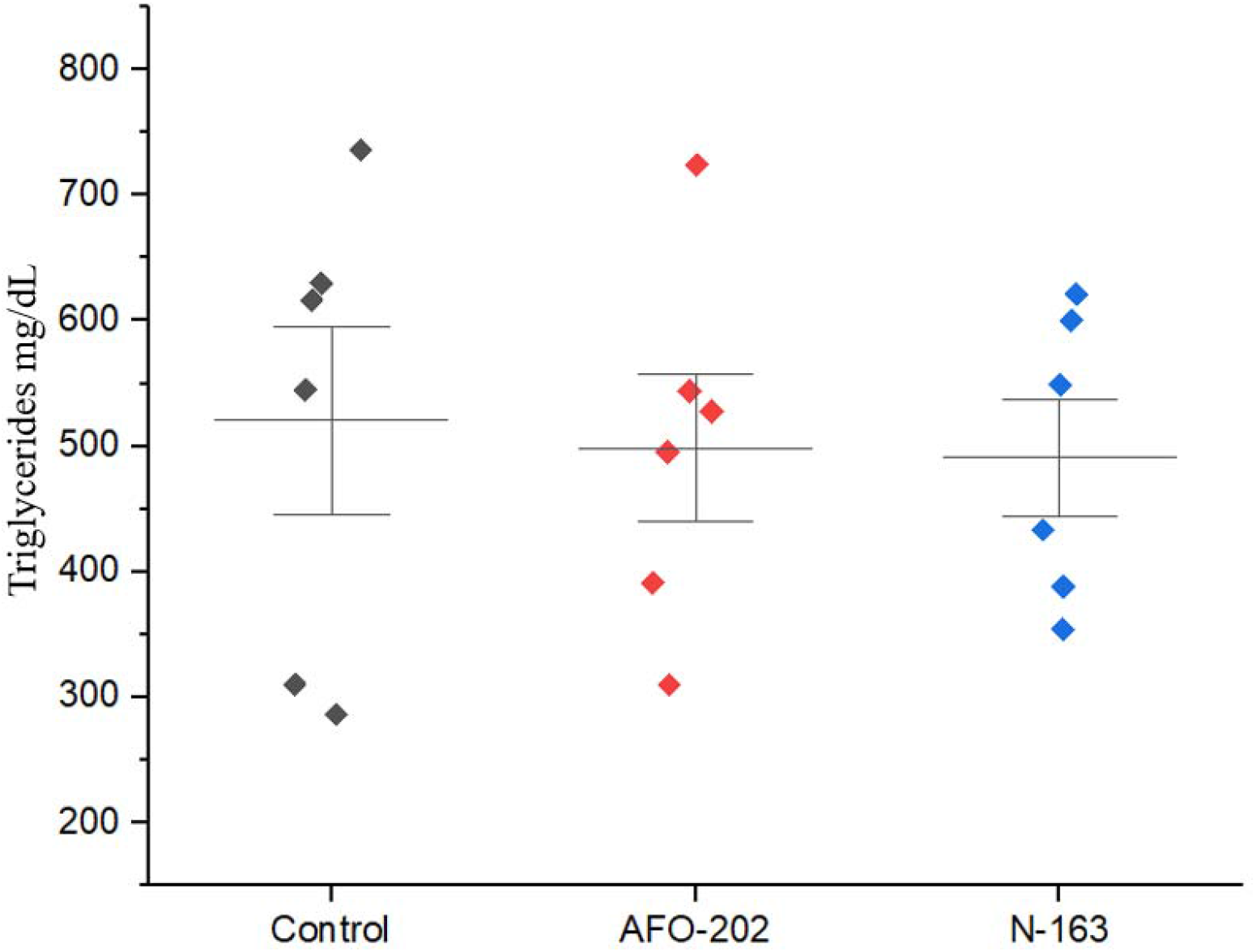
Triglyceride levels in male KK-Ay mice administered solution orally for 28 days showing marginal reduction in Group 3 (N-163 beta glucan) compared to Groups 2 (AFO-202 beta glucan) and 1 (control)

## Discussion

Dyslipidaemia is an emerging epidemic worldwide, and nearly 39% of the global population has elevated cholesterol levels. Dyslipidaemia is the cause of 26.9% of morbidity and 28% of mortality, according to the Global Burden of Diseases, Injuries, and Risk Factors study [11]. The liver plays a major role in the metabolic pathways of glucose and lipids. In particular, NEFA has been indicated as a major cause for the progression of non-alcoholic liver injury because the liver is responsible for taking up serum FFA and manufacturing, storing and transporting lipid metabolites. The circulating NEFA pool contributes to the majority of the FFA that flow to the liver; hence, elevated NEFA contributes to fatty liver disease progression [12]. In addition, several reports have pointed out the key role of chronic inflammation in the pathogenesis of obesity-related metabolic dysfunction and non-alcoholic fatty liver disease (NAFLD) [13]. This inflammatory pathway, caused by a high-fat diet and elevated lipid levels, leads to insulin resistance and the progression of metabolic syndromes and NAFLD [14]. Adipose tissue is a potential source of elevated circulating inflammatory cytokines such as TNF-α and IL-6, especially with the increased numbers of macrophages that accumulate in adipose tissue in obese individuals contributing to the secretion of these cytokines [15]. NEFA induces the macrophages to secrete inflammatory cytokines [15]. Thus, the entire cascade of metabolic syndrome, diabetes, obesity and an inflammatory state contributing to cancer can be attributed to the lipotoxicity and may benefit from strategies that target lipids, especially NEFA level normalization.

Beta glucans, especially derived from A. *pullulans*, are potent immunomodulators that have already been shown to decrease IL-6 and TNF-α levels, in addition to positively regulating Akt/PI3K and peroxisome proliferator-activated receptor γ (PPARγ) signalling pathways, which are principal regulators of adipogenesis and glucose metabolism [16]. These beta glucans are capable of modulating the immune response by suppressing inflammatory cytokines [17] without over-activation. The beta glucans are capable of eliciting a unique immune response, binding directly with immune cells such as macrophages and, importantly, downregulating the abnormal macrophages while activating the normal macrophages [18-20]. Therefore, beta glucans are considered a potential strategy to counter the ill effects of dyslipidaemia and reduce elevated NEFA levels.

In the current study, N-163 beta glucan showed beneficial reductions of NEFA and triglyceride levels, as compared to the AFO-202 beta glucan and the control. At the end of the treatment period, the blood glucose levels in Groups 2 and 3 were within the range of 588 to 615 mg/dL. In the literature (KK-Ay/Ta Jcl Mouse Data Collection, Nihon Clare Co., Ltd., 1994), the blood glucose level in 10-week-old KK-Ay male mice was reported to be 333 ± 33 mg/dL when not fasting. This suggests that the present experiment reproduced a hyperglycaemic state similar to or higher than those found in the literature throughout the treatment period, thus recapitulating the metabolic syndrome that occurs in humans.

One major limitation of the study is that the dosage was based on human consumption levels of beta glucans, and the earlier reports [8,9] have indicated the normalization of lipid levels at least 2 months after the treatment. Because the present study lasted only 28 days, a dose-escalation study will be more useful in the context of studying the effects of beta glucans in a shorter time duration. Further evaluation of these two beta glucans in balancing the parameters of relevance to metabolic syndrome, glucotoxicity and lipotoxicity and ensuring control of the inflammatory cascade in both healthy and diseased patients may shed more light on their efficacy and probable mechanisms.

## Conclusion

Repeated oral administration of beta glucans derived from AFO-202 and N-163 strains of *A. pullulans* for 28 days resulted in lower triglyceride and NEFA levels among male KK-Ay mice than before the start of treatment. In particular, the NEFA levels in N-163-fed mice were lower than those in the AFO-202 group, making further research recommended on anti-inflammatory efficacy, which may help with designing effective therapeutic strategies for treating and preventing fibrotic diseases that develop due to dyslipidaemia-related cascade.

## Ethics Approval

The protocol approval was obtained by the ethics committee of Toya Laboratory, HOKUDO Co (Ref no: HKD47047). The study was conducted in accordance with the HOKUDO Animal Experiment Regulations following the Act on Welfare and Management of Animals (Ministry of the Environment, Japan, Act No. 105 of October 1, 1973), standards relating to the care and management of laboratory animals and relief of pain (Notice No.88 of the Ministry of the Environment, Japan, April 28, 2006) and the guidelines for proper conduct of animal experiments (Science Council of Japan, June 1, 2006).

## Availability of data and material

All data generated or analysed during this study are included in this manuscript

## Acknowledgements

The authors thank

a. Mr. Yoshio Morozumi, Ms. Yoshiko Amikura of GN Corporation, Japan for their liaison assistance with the conduct of the study.
b. Loyola-ICAM College of Engineering and Technology (LICET) for their support to our research work.

## Notes

### Competing Interest Statement

1.Author Samuel Abraham is a shareholder in GN Corporation, Japan which in turn is a shareholder in the manufacturing company of the Beta Glucans described in the study.
2.Author Takashi Onaka is a shareholder and Yasunori Ikeue is member of the board in the manufacturing company of the Beta Glucans described in the study.

